# Systematic identification and characterization of *Aedes aegypti* long noncoding RNAs

**DOI:** 10.1101/422170

**Authors:** Azali Azlan, Sattam M. Obeidat, Muhammad Amir Yunus, Ghows Azzam

## Abstract

Long noncoding RNAs (lncRNAs) play diverse roles in biological process including developmental regulation and host-pathogen interactions. *Aedes aegypti* (*Ae. aegypti*), a blood-sucking mosquito, is the principal vector responsible for replication and transmission of arboviruses including dengue, zika, and chikungunya virus. Systematic identification and developmental characterisation of *Ae. aegypti* lncRNAs are still limited. We performed genome-wide identification of lncRNAs followed by developmental profiling of lncRNA expression in *Ae. aegypti.* We identified a total of 4,689 novel lncRNA transcripts, of which 2,064, 2,076, and 549 were intergenic, intronic, and antisense respectively. *Ae. aegypti* lncRNAs shared many of the characteristics with other species including low expression, low GC content, short in length, low conservation, and their expression tended to be correlated with neighbouring and antisense protein-coding genes. Subsets of lncRNAs showed evidence that they were maternally inherited, suggesting potential roles in early-stage embryos. Additionally, lncRNAs showed higher tendency to be expressed in developmental and temporal specific manner. Upon infection of *Ae. aegypti* cells with dengue virus serotype 1, we identified 2,335 differentially expressed transcripts, 957 of which were lncRNA transcripts. The systematic annotation, developmental profiling, and transcriptional response upon virus infection provide foundation for future investigation on the function of *Ae. aegypti* lncRNAs.

## Introduction

*Aedes aegypti* (*Ae. aegypti*), a blood-sucking mosquito, is the principal vector responsible for replication and transmission of arboviruses such as dengue (DENV), Chikugunya (CHIKV), and Zika (ZIKV) virus. Although virus infections of the vertebrate host are acute and often associated with disease, the maintenance of these pathogens in nature generally requires the establishment of a persistent non-lethal infection in the insect host. This is also true in the context of dengue virus infection. While dengue-infected humans show serious symptoms of illnesses, interestingly, infected *Ae. aegypti* mosquitoes are generally asymptomatic, suggesting that these vectors possess intrinsic antiviral immunity to resist or tolerate the virus infection. A number of studies have shown that, upon flavivirus infection, major transcriptomic changes occurred within *Ae. aegypti.* Such changes involved a wide range of host genes including genes related to immunity, metabolic pathways, and trafficking. Although protein-coding genes have been the central focus, previous report indicated that *Ae. aegypti* lncRNAs were also involved in virus-host interaction^1,2^.

Advances in high-throughput sequencing technologies coupled with bioinformatics analyses have enabled the identification of a significant number of lncRNAs in various species. Such lncRNAs exert their functions by a range of mechanisms. For instance, several lncRNAs regulate gene expression by sequestering miRNAs and transcription factors ^3,4^. Other lncRNAs have been shown to regulate the activity of chromatin modifying proteins; hence, modulating gene expression. In *Ae. aegypti*, it had been reported that lncRNA was involved in virus-host interaction. Knockdown of lincRNA_1317 in *Ae. aegypti* cells caused higher replication of DENV^1^. Although lncRNAs have been systematically identified in many organisms, most lncRNAs have not been functionally characterised.

Recently, the latest version of *Ae. aegypti* genome (AaegL5) was released, and the assembly was up to chromosome-length scaffolds, which is more contiguous than the previous AaegL3 and AaegL4 assemblies^5^. This prompted us to perform novel lncRNA identification and characterisation using the latest genome release as reference. Previous study^1^ reported that a total of 3,482 intergenic lncRNA (lincRNA) was identified in *Ae. aegypti*. However, the identification was performed using previous version of *Ae. aegypti* genome (AaegL3), and only lncRNAs residing in intergenic region were annotated.

Here, we report genome-wide identification and characterisation of lncRNAs in *Ae. aegypti.* Identification of lncRNAs was performed using a combination of RNA-seq based *de novo* transcript discovery and rigorous filtering of putative protein-coding transcripts. In this study, we defined a high-confidence set of 4,689 novel lncRNA transcripts −2,064 2,076, and 549 were intergenic, intronic, and antisense respectively. We then characterised the newly identified lncRNAs by diverse features such as transcript structures, conservation, and developmental expression. Previous studies have reported global transcriptional responses of lncRNAs upon infection with dengue virus serotype 2 (DENV2) and ZIKV. Here, we examined the transcriptome profiles in *Ae. aegypti* cell (Aag2) in response to DENV serotype 1 (DENV1) infection. Collectively, genome-wide annotation and characterisation of *Ae. aegypti* lncRNAs provide valuable resources for future genetics and molecular studies in this harmful mosquito vector.

### Identification of lncRNA

To perform lncRNA prediction, we used a total of 117 datasets of Illumina high-throughput sequencing derived from *Ae. aegypti* mosquito and Aag2 cell, a widely used *Ae. aegypti* derived cell line^6^. An overview of lncRNA identification pipeline can be found in Fig 1. The pipeline developed in this study was adapted with few modifications from previous reports ^1,7,8^. Briefly, each dataset (both paired-end and single-end) was individually mapped using HISAT2^9^. The resulting alignment files were used for transcriptome assembly, and the assemblies were merged into a single unified transcriptome. Both transcriptome assembly and merging were performed using Stringtie^10^. Then, we used Gffcompare to annotate and compare the unified transcriptome assembly with reference annotation (AaegL5.1, VectorBase). We classified lncRNA transcripts based on their position relative to annotated genes derived from AaegL5.1 assembly (VectorBase). We only selected transcripts with class code “i”, “u”, and “x” that denote intronic, intergenic, and antisense to reference genes for downstream analysis.

**Fig 1.**
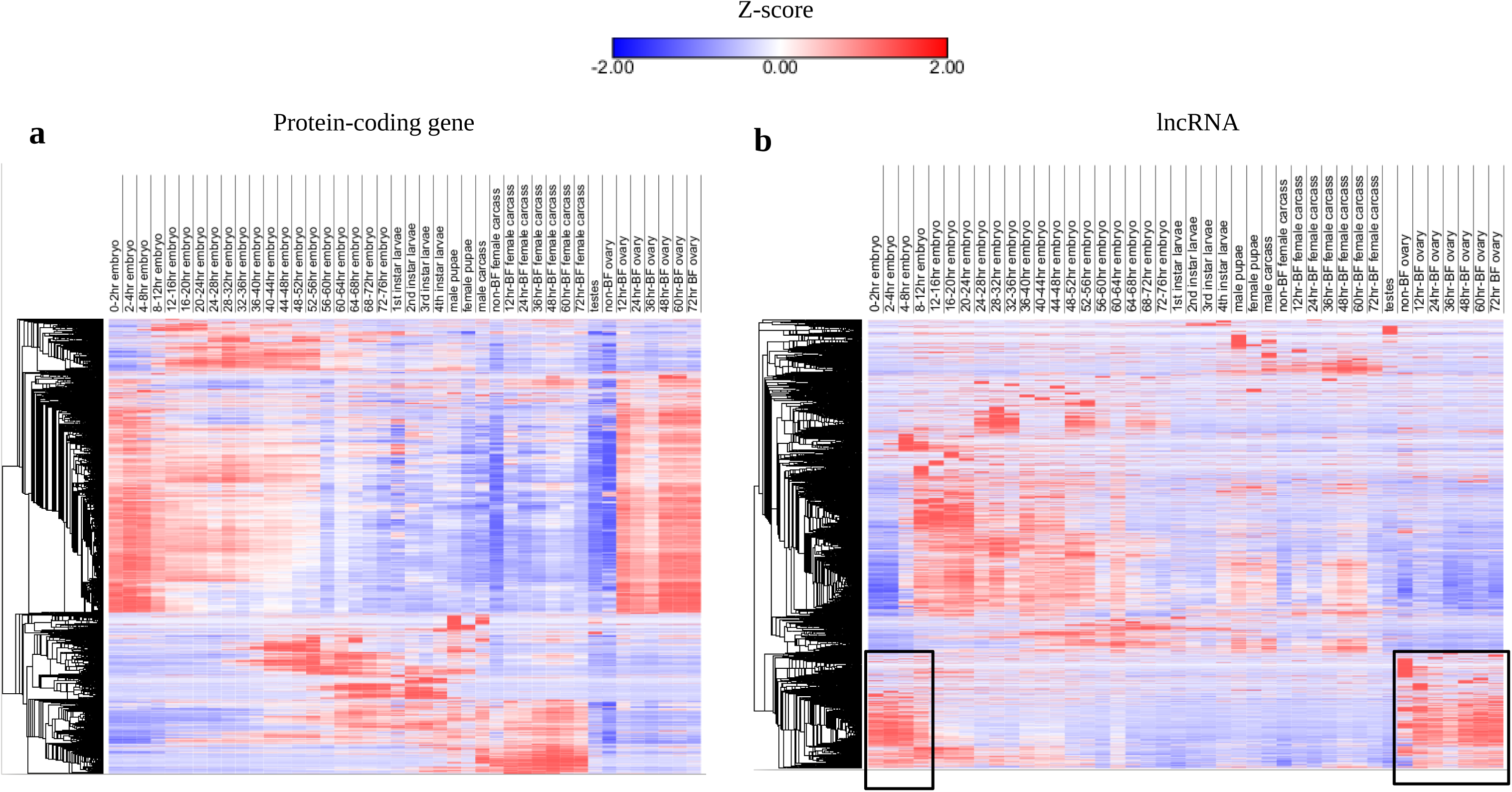
lncRNA identification. Overview of lncRNA identification pipeline.

To get confident lncRNA transcripts, we performed multiple steps of filtering transcripts having coding potential or open-reading frame (Fig 1). The steps involved TransDecoder^11^, CPAT^12^ and finally BLASTX. We then removed lncRNA candidates that did not have strand information. Detailed description on the prediction analysis and parameter used can be found in Material and Methods section. Using this pipeline, we identified a set of 4,689 novel lncRNA transcript isoforms derived from 3,621 loci. Of these 4,689 lncRNA transcripts, 2,064 and 2,076 were intergenic and intronic respectively, while the remaining 549 transcripts were antisense to reference genes. Currently, AaegL5.1 annotation catalogs 4,155 lncRNA transcripts. Here, we provided another set of 4,689 lncRNA transcripts, making a total of 8,844 lncRNAs in *Ae. aegypti*.

### Characterisation of *Ae. aegypti* lncRNA

To examine whether lncRNAs identified in this study exhibit typical characteristics observed in other species^8,13–15^, we analysed features such as coding potential, sequence length, GC content and sequence conservation with closely related species. Since lncRNAs are strictly defined by their inability to code for protein, we determined coding probability of our newly identified lncRNAs and compared them with known lncRNA, 3’UTR, 5’UTR, and protein-coding mRNA. We discovered that, similar to other non-coding sequence such as known lncRNA, 3’UTR, and 5’UTR, our novel lncRNA transcripts have extremely low coding probability when compared to protein-coding mRNA (Fig 2a). Beside that, we found that both novel and known lncRNAs (provided by AaegL5.1 annotation) were shorter than protein-coding transcripts (Fig 2b). Mean length of novel and known lncRNAs was 825.4 bp and 745.6 bp respectively, while protein-coding mRNA has an average length of 3330 bp.

**Fig 2.**
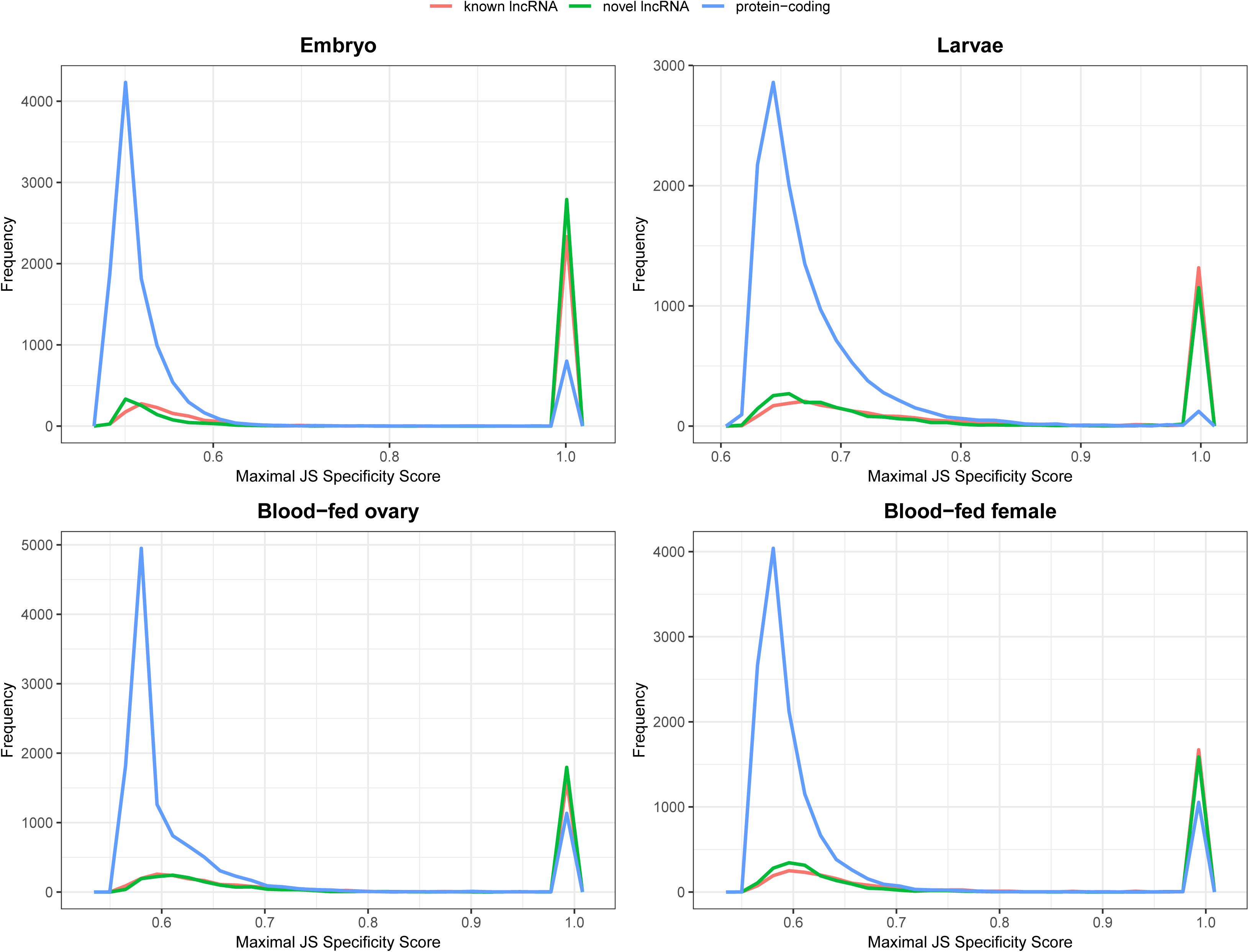
Characterization of *Ae aegypti* lncRNA. (a) Coding probability of lncRNA, 3’UTR, 5’UTR and mRNA. (b) Sequence length distribution of lncRNA and mRNA. (c) GC content. (d) Repeat content

Similar to previous reports^1,8^, we observed that lncRNAs identified in this study had slightly lower GC content compared to protein-coding mRNAs (Fig 2c). For instances, mean GC content of novel lncRNA and mRNA was 40.1% and 46.4% respectively. Known lncRNA, on the other hand, had relatively similar mean GC content to novel lncRNA (40.8%), while average GC of 5’UTR and 3’UTR sequence was 43.1% and 34.6% respectively. Overall, GC content of non-coding sequence was relatively lower than coding sequence.

To determine the conservation level of *Ae. aegypti* lncRNAs, we performed BLASTN against other insect genomes including *Ae. albopictus, C, quinquifasciatus*, *An, gambiae* and *D. melanogaster*, Bit score derived from BLAST algorithm was used to determine the level of sequence similarity of *Ae. aegypti* lncRNAs with previously mentioned insect genomes^1^. Similarly, we also performed BLASTN of *Ae. aegypti* protein-coding mRNA for comparison. We discovered that both lncRNAs and protein-coding mRNAs displayed high degree of sequence similarity with *Ae. albopictus*, suggesting that they were presumably genus specific (Supplementary Fig. 1). In general, compared to protein-coding mRNA, *Ae. aegypti* lncRNAs exhibited lower sequence conservation.

It was reported that, in contrast to coding gene, lncRNAs harbour higher composition of repeat elements^16,7^. To test if similar occurrence held true in *Ae. aegypti*, we determined the fraction of repeat element in the exons of lncRNAs and protein-coding mRNAs. As expected, we discovered that more than 20% of nt of novel lncRNAs were made up of repeat elements (Fig 2d). Meanwhile, 6% and 1% of known lncRNAs and protein-coding mRNAs were composed with repeats. Taken together, *Ae. aegypti* lncRNAs shared many characteristics with other species: relatively short in length, relatively lower GC content, and higher repeat content.

### Developmental expression of *Ae. aegypti* lncRNAs

To examine the developmental expression of *Ae. aegypti* lncRNAs, we analysed public dataset (SRP026319) which provided RNA-seq data of specific developmental stages, ranging from early embryo up to adult mosquitoes. These developmental stages include specific time interval of embryonic development, larval stages, sex-biased expression of male versus female pupae and carcass, blood-fed versus nonblood-fed ovary and female carcass, and testes-specific expression. Similar to protein-coding genes, *Ae. aegypti* lncRNA genes exhibited stage-specific (embryo, larva, and pupal stages), sex-specific, and blood-fed versus nonblood-fed (ovary and female carcass) expression pattern (Fig 3). In addition, each time point in the development had distinct lncRNA expression pattern. For instance, there was a subset of lncRNAs that were highly enriched during early embryonic development (0-8 hour embryo) as compared to later time points (8-76 hour embryo). A total of 1,848 lncRNA genes consisting of 24.7% of the total expressed lncRNAs were specifically highly expressed at this early embryonic stage. Interestingly, these early embryo-specific lncRNAs were also highly expressed in blood-fed ovary, suggesting that these lncRNAs may have potentially maternally inherited and possess essential roles in early development.

**Fig 3.**
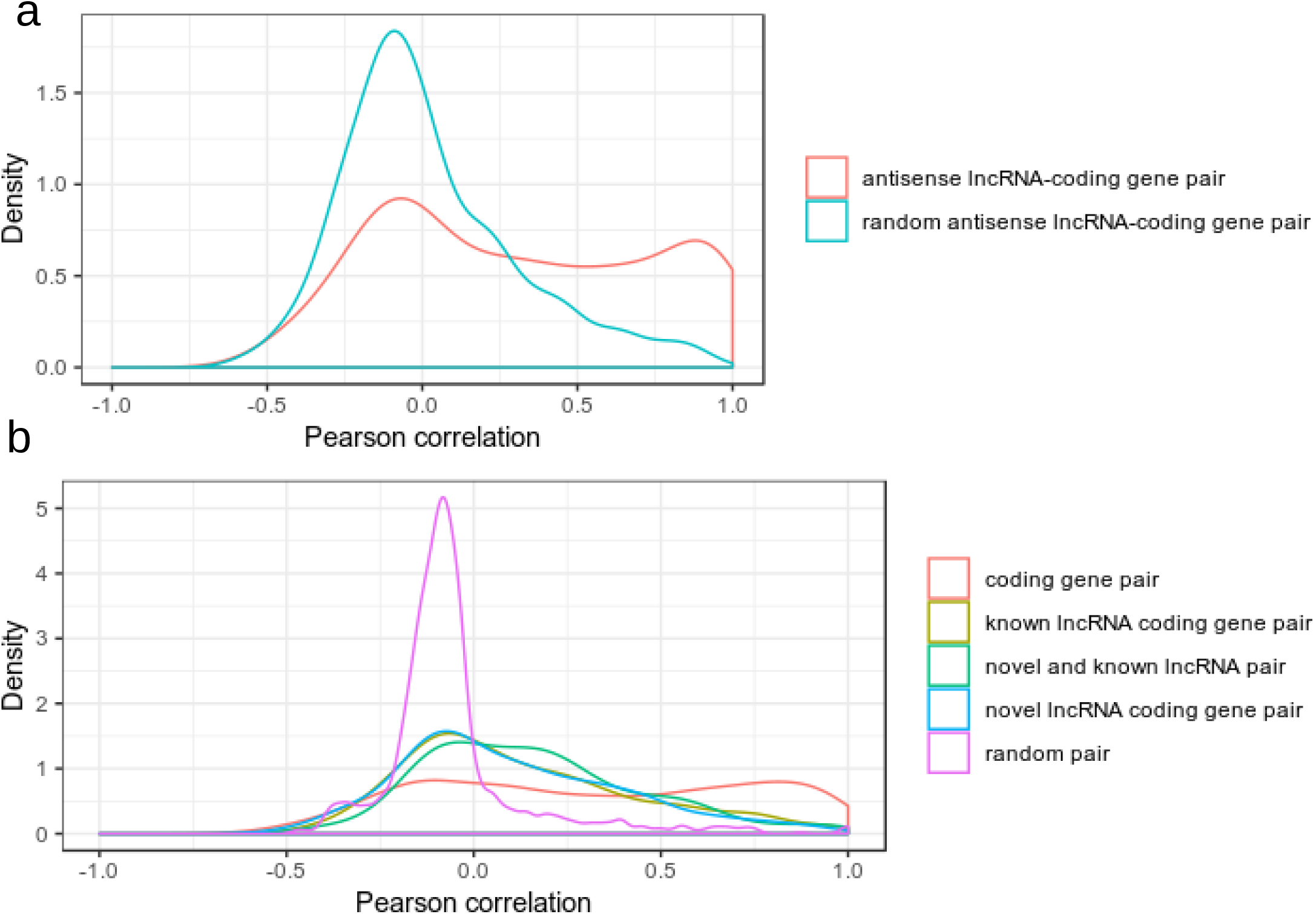
Hierarchical clustering of protein-coding and lncRNA expression. Hierarchical clustering of protein-coding (a) and lncRNA (b) across developmental stages. Hierarchical clustering analysis was done in Morpheus (https://software.broadinstitute.org/morpheus) based on Pearson correlation of z scores of lncRNA and protein-coding genes. Boxes in (b) indicate a subset of lncRNAs that are presumably maternally inherited.

Previous report showed that lncRNAs displayed a more temporally specific expression pattern than protein-coding genes^17^. To test whether or not this is true for *Ae. aegypti* lncRNAs, we computed specificity score of each lncRNA using Jensen-Shannon (JS) score^18,17^. The JS score ranged from 0 to 1, with 1 indicated perfect specificity. Here, we computed JS score across two stages namely embryo and larvae, and two conditions which were blood-fed ovary and female carcass, all of which were sampled in a timely fashion. In all four time-course samples namely embryo, larvae, blood-fed female carcass and ovary, we discovered that novel and known lncRNAs had higher JS specificity score in all four samples (Fig. 4). Meanwhile, fraction of protein-coding genes having JS score of 1in all four samples were lower than lncRNAs (Fig. 4). Although lncRNAs had higher developmental and temporal specificity, across all four time-course samples being analysed, the overall expression of lncRNAs was lower than that of protein-coding genes (Supplementary Fig. 2).

**Fig 4.**
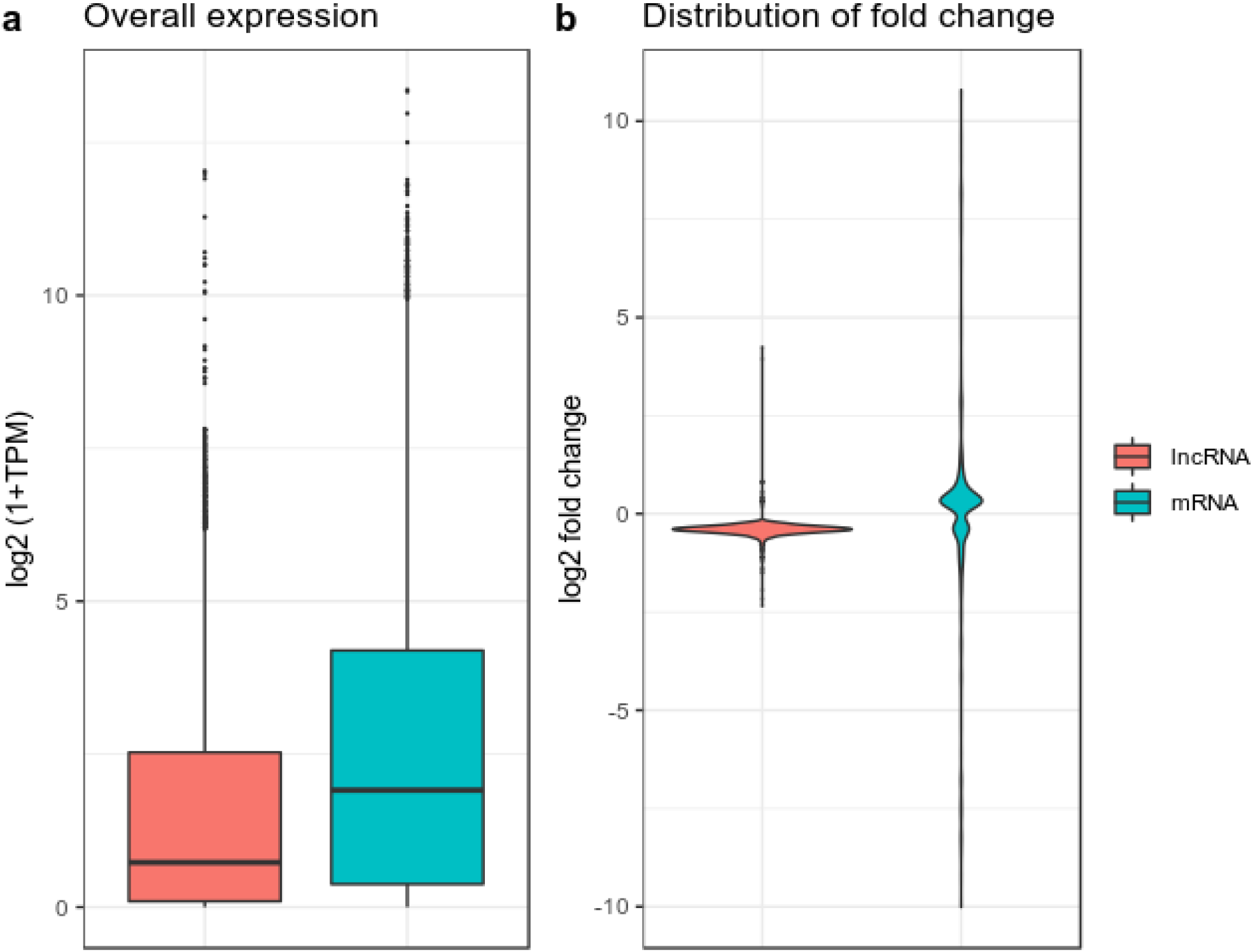
Maximal JS specificity score

Previous studies have shown that lncRNAs are favourably resided in close proximity to genes with developmental functions^19,20^. In addition, this close physical proximity of lncRNAs and protein-coding genes resulted in their expression to be strongly correlated. We asked whether *Ae. aegypti* lncRNAs showed correlation in expression between neighboring protein-coding genes. We also examined the correlation in expression of lncRNAs that were antisense to protein-coding genes. By analysing the expression of 42 developmental samples (Fig. 3), we discovered that fraction of positively correlated (Pearson correlation, p-value < 0.05) antisense lncRNA-coding gene pair was higher than that of randomly assigned antisense lncRNA-coding gene pair (Fig 5). Analysis of neighboring genes (within 10 kb) revealed that the expression of lncRNAs and their nearest neighboring genes showed a slightly higher degree of correlation compared to random gene pairs. In both cases of random neighbouring pair and random antisense lncRNA-coding gene pair, the majority of the pairs had correlation near to zero value (Fig 5).

**Fig 5.**
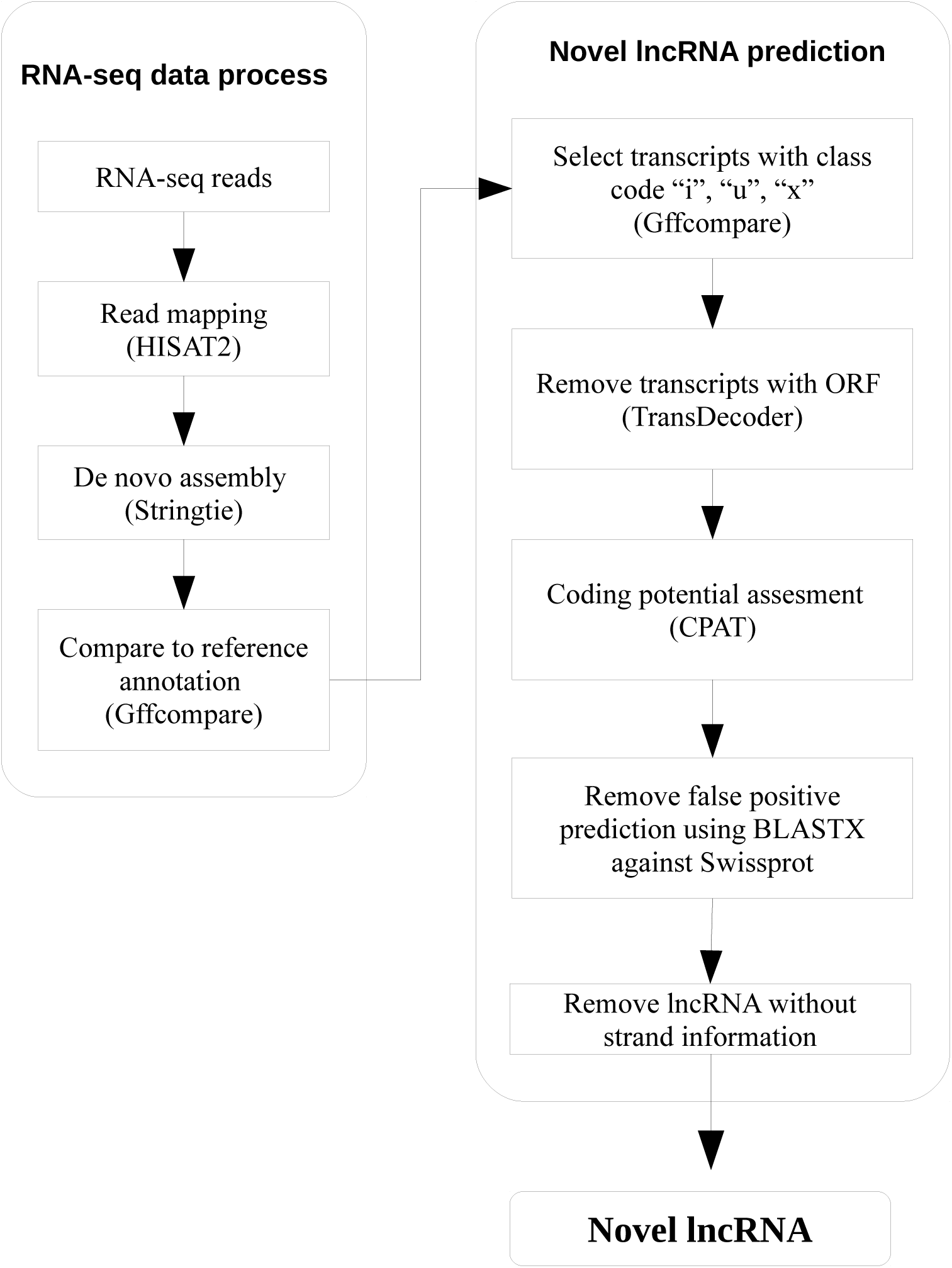
Correlation of expression analysis. (a) Pearson correlation of antisense lncRNA expression with its corresponding genes compared to that of antisense lncRNA pair with random protein-coding gene (b) Pearson correlation of gene expression within 10 kb from each other

### *Ae. aegypti* lncRNAs were differentially expressed following DENV1 infection

Previous studies reported that lncRNAs in *Ae. aegypti* underwent differential expression upon dengue virus serotype 2 (DENV2) and ZIKV infection. In this study, we chose another DENV serotype, namely serotype 1, and observed the transcriptional response of *Ae. aegypti* lncRNAs upon infection. Therefore, we generated paired-end RNA-seq libraries derived from triplicates of Aag2 cells infected with DENV1. To avoid bias in library size which affects fold change calculation, we performed differential expression analysis of protein-coding genes and lncRNAs together in a single transcriptome reference using Salmon v0.9^21^ followed by edgeR^22^. Here, differential expression analysis was performed at transcript level. A total of 2,335 transcripts were differentially expressed (P-value < 0.05) upon DENV1 infection, 957 of which were lncRNA transcripts.

In general, we found that across all six samples (3 uninfected and 3 DENV1-infected samples), expression of lncRNA was lower than protein-coding mRNAs (Fig 6a). Besides, distribution of fold change (FC) of differentially expressed lncRNAs (P-value < 0.05) were much lower than that of mRNAs (Fig 6b). This observation suggests that unlike mRNA, lncRNA expression was less responsive to viral infection. For example, log2 FC of mRNA can extend to more than 10 fold in either direction (−10 and 10 fold), but in lncRNA, log2 FC ranged from −2 to 5. Besides, most of the differentially expressed lncRNAs (953 out of 957) transcripts had log2 FC of less than 2 in either direction.

**Fig 6.**
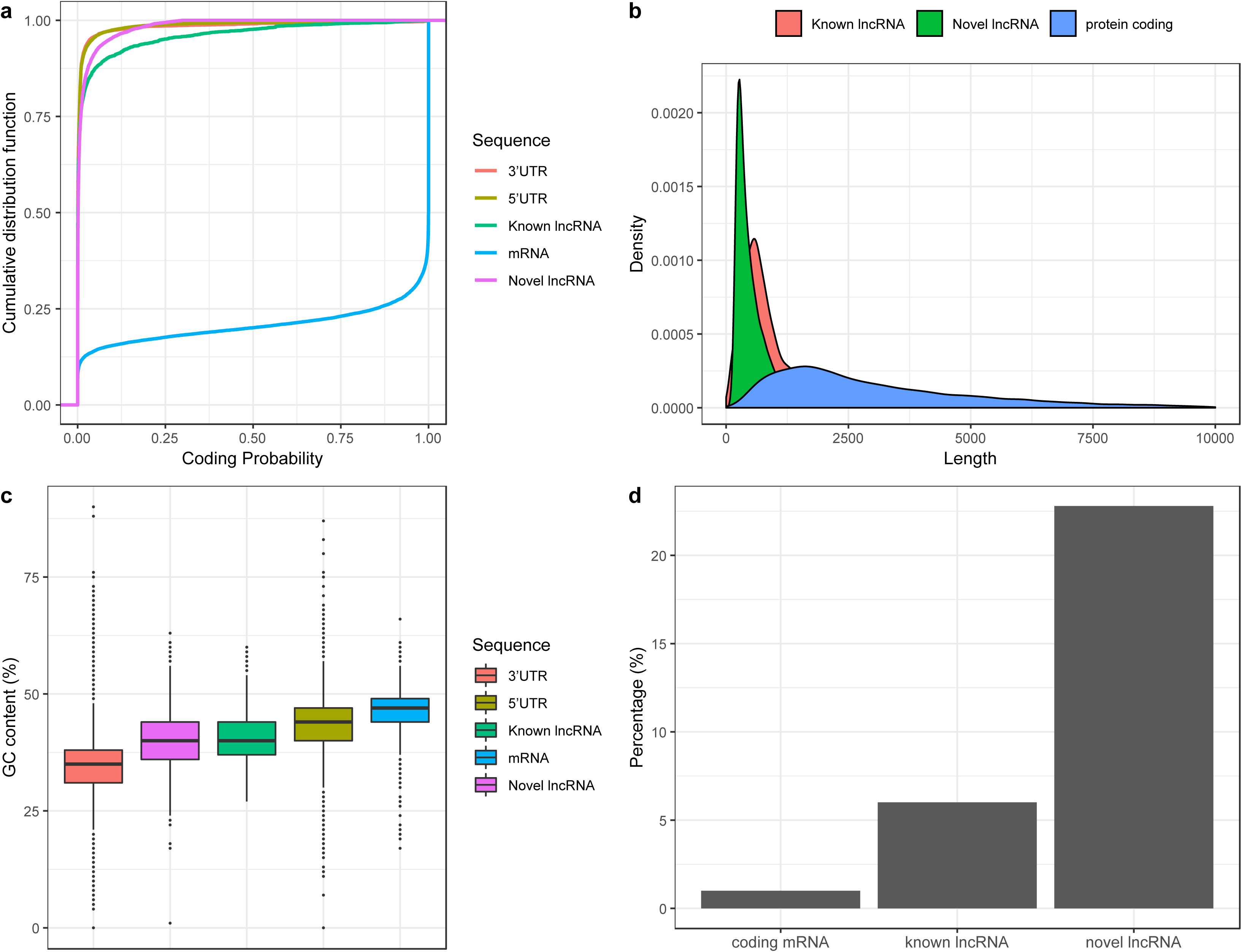
Overall distribution of expression and fold change of lncRNA and mRNA upon DENV1 infection

## Discussion

The field of lncRNA has become increasingly important in many areas of biology particularly infectious disease, immunity, and pathogenesis^1,8,14,23^. High-throughput sequencing combined with bioinformatics enable scientists to uncover comprehensive repertoire of lncRNA in many species. Here, we present a comprehensive lncRNA annotation using the latest genome reference of *Ae. aegypti* (AaegL5). Due to the recent release of *Ae. aegypti* genome (AaegL5) equipped with improved gene set annotation^5^, we decided to perform lncRNA identification using this latest genome reference. Unlike previous annotation that mainly focused on *Ae. aegypti* intergenic lncRNA, here, we also annotated lncRNAs residing within the introns and antisense to reference genes. We used large RNA-seq datasets for lncRNA annotation; hence, enabling identification of *Ae. aegypti* lncRNAs that are lowly expressed.

In addition, we used recently developed and improved softwares including HISAT^9^, StringTie^10^, and Salmon^21^ for novel lncRNA prediction and expression analysis. HISAT is considered to be faster with equal or better accuracy than other spliced aligner methods such as Tophat^24^ and STAR^25^. Meanwhile, compared to other transcript assembly software such as Cufflinks^26^ and Scripture^27^, StringTie has been shown to produce more comprehensive and accurate transcriptome reconstruction from RNA-seq data^10^. Salmon was used for transcript abundance quantification in this study due to its rapidness and accuracy since the algorithm is able to correct for fragment GC content bias^21^. Taken together, the use of large RNA-seq datasets, the most recent genome reference, and improved algorithms in bioinformatics analyses enable us to provide, to date, the most comprehensive and confident lncRNA annotation in *Ae. aegypti* transcriptome.

Similar to previous reports^1,8,17,23,28^, we discovered that lncRNAs identified in our study exhibited typical characteristics of lncRNAs found in other species including vertebrates^7^. Such characteristics are lower GC content, shorter in length, and low sequence conservation even among closely related species. Analysis of *Ae. aegypti* developmental expression revealed that the expression of lncRNAs is highly temporally specific relative to that of coding genes. In other words, lncRNAs have a much narrower time window in expression than coding genes. Such specificity in expression suggests essential roles of lncRNAs in *Ae. aegypti*, especially in regulating the timing of developmental transition.

Several studies have pointed out that lncRNAs are essential in development and cell differentiation^29^. For example, lncRNAs emerge as important regulators of embryogenesis, by means of directing pluripotency and differentiation^30^ In this study, we provided an evidence of maternal inheritance of *Ae. aegypti* lncRNAs. This further corroborated previous findings that highlighted the importance of lncRNAs in embryogenesis^20,17,31^ We observed a fraction of lncRNAs that was highly expressed in blood-fed ovary, and the expression persisted up to 8-12 hour embryonic stage. This narrow time window in early embryo was related to maternal-zygotic transition stage^32^. Since transcription from the zygotic genome has not been activated during early embryonic stage, these highly expressed lncRNAs must be maternally provided; suggesting that they might play roles in basic biosynthesis processes in the early embryo, specification of initial cell fate and pattern formation. Meanwhile, lncRNAs expressed at later embryonic, larval and pupal stages are putatively responsible in organogenesis.

Upon DENV1 infection, we observed that both protein-coding and lncRNAs experienced overall changes in expression level. However, the overall level of fold change displayed by lncRNAs was not as high as protein-coding genes. This observation however, did not necessarily negate the possibility of lncRNAs having potential roles in immunity. For example, RNAi-mediated knockdown of one lncRNA candidate (lincRNA_1317) in *Ae. aegypti* resulted in higher viral replication^1^. Therefore, intensive investigation using loss or gain of function approach on lncRNAs is crucial in dissecting their functions in mosquito-virus interaction.

In summary, we provided a comprehensive list of lncRNAs in *Ae. aegypti* using the latest genome version. This study highlights numerous roles of lncRNAs in *Ae. aegypti* especially in development and virus-host interaction.

## Materials and Methods

### Cell culture and virus

*Ae. aegypti* Aag2 cell and *Ae. albopictus* C6/36 cell were cultured in Leibovitz’s L-15 medium (Gibco, 41300039), supplemented with 10% Fetal Bovine Serum (FBS, Gibco, 10270) and 10% Tryptose Phosphate Broth (TPB) solution (Sigma, T9157). Both Aag2 and C6/36 cells were incubated at 25ºC without CO_2_. Vero E6 cells were cultured at 37ºC in Dulbecco’s modified Eagles Medium (DMEM, Gibco, 11995065) supplemented with 10% FBS (Gibco, 10270), and 5% CO_2_. DENV1, Hawaiian strain was propagated in C6/36 cells and titered using Vero E6 cells. Determination of DENV1 titer was done using 50% tissue culture infectious dose – cytopathic effect (TCID50-CPE) as previously described^33^. DENV1 used in this study was a gift from Dr. David Perera, University Malaysia Sarawak. Aag2 cell line was a kind gift from Dr. Ronald P. Van Rij from Radbound Institute for Molecular Life Sciences, Netherlands.

### Virus infection, RNA isolation and sequencing

Aag2 cells were infected with DENV1 at multiplicity of infection (MOI) of 0.5. At 72 hours post infection (hpi), the culture medium was removed, and RNA isolation was carried out using miRNeasy Mini Kit 50 (Qiagen, 217004) according to the manufacturer’s protocol. Total RNA was then subjected to next-generation sequencing. The RNA-sequencing libraries were prepared using standard Illumina protocols and sequenced using HiSeq2500 platform generating paired-end reads of 150 in size.

### Verification of DENV1 infection

Total RNA of both uninfected and DENV1-infected samples were subjectred to cDNA synthesis using Tetro cDNA synthesis kit (Bioline, BIO-65042). Reverse primer for DENV1 was included in the reaction set-up. This followed with PCR and gel electrophoresis using forward and reverse primers of DENV1. Primers used for DENV1 verification were taken from previous study^34^.

### RNA-seq data preparation

RNA-seq datasets generated in this study (Aag2 cell infected with DENV1) are deposited in the Short Read Archive database under the accession code: PRJNA509977. Publicly available datasets were downloaded from NCBI Sequence Reads Archive (SRA) and ArrayExpress with accession numbers SRP041845, SRP047470, SRP046160, SRP115939, E-MTAB-1635, SRP035216, SRP065731, SRP065119, SRA048559, PRJEB13078, SRP026319^2,35–40^. Adapters were removed using Trimmomatic version 0.38^41^, and reads with average quality score (Phred Score) above 20 were retained for downstream analysis.

### lncRNA identification

Each library (both paired-end and single-end) was individually mapped against *Ae. aegypti* genome (AaegL5) using HISAT2 version 2.1.0^9^. The resulting alignment files were used as input for transcript assembly using Stringtie version 1.3.2^10^. We used reference annotation file of AaegL5 (VectorBase) to guide the assembly only without discarding the assembly of novel transcripts. Here, we set the minimum assembled transcript length to be 200 bp. Then, the output gtf files were merged into a single unified transcriptome using Stringtie merge^10^. Only input transcripts of more than 1 FPKM and TPM were included in the merging. Then, we compared the assembled unified transcript to a reference annotation of AaegL5 (VectorBase) using Gffcompare (https://github.com/gpertea/gffcompare). For the purpose of lncRNA prediction, we only retained transcripts with class code “i”, “u”, and “x”. The transcripts were then subjected to coding potential prediction. We used TransDecoder^11^ to identify transcripts having open-reading frame (ORF), and those having ORF were discarded. The remaining transcripts were then subjected to a coding potential assessment toll (CPAT)^12^. We set the same cut-off as previous study in *Ae. aegypti* which is less than 0.3^1^. Transcripts having coding potential more than 0.3 were discarded. To exclude false positive prediction, we used BLASTX against Swissprot database, and transcripts having E-value of less than 10^-5^ were removed.

### Identification of differentially expressed transcripts

We used Salmon version 0.10.1^21^ to quantify the expression of transcripts. Read count for each transcript was subjected for differential analysis using edgeR^22^ in R/Bioconductor environment.

### Quantitative real-time PCR (qRT-PCR)

cDNA synthesis was done using QuantiNova Reverse Transcription Kit (Qiagen) followed by qRT-PCR with QuantiNova SYBR Green PCR kit (Qiagen) according to manufacturer’s protocol. *RPS17* was used as housekeeping gene^42^, and the experiment was performed using Applied Biosystems Step One PlusTM Real-Time PCR System.

## List of Supporting Information

**S1 Table. Genomic coordinates of lncRNA candidates identified in *Ae. aegypti***

**S2 Table. List of differentially expressed lncRNA S3 Table. List of primers used in this study**

**S1 Fig. Similarity bit score of lncRNA and mRNA with closely related insect genomes**

**S2 Fig. Distribution of lncRNA and coding gene expression in embryo, larvae, blood-fed ovary and female carcass**

**S3 Fig. Validation of differentially expressed lncRNAs by qRT-PCR**

Two-tailed *t-*test was used for all comparisons; **p < 0.05, ***p < 0.01. Expression levels were normalised to uninfected samples. Error bars indicate standard error of the mean (SEM) of three independent experiments.

