## Supplementary figures and images for "Systematic identification and characterization of *Aedes aegypti* long noncoding RNAs"

### Supplementary Figure S1

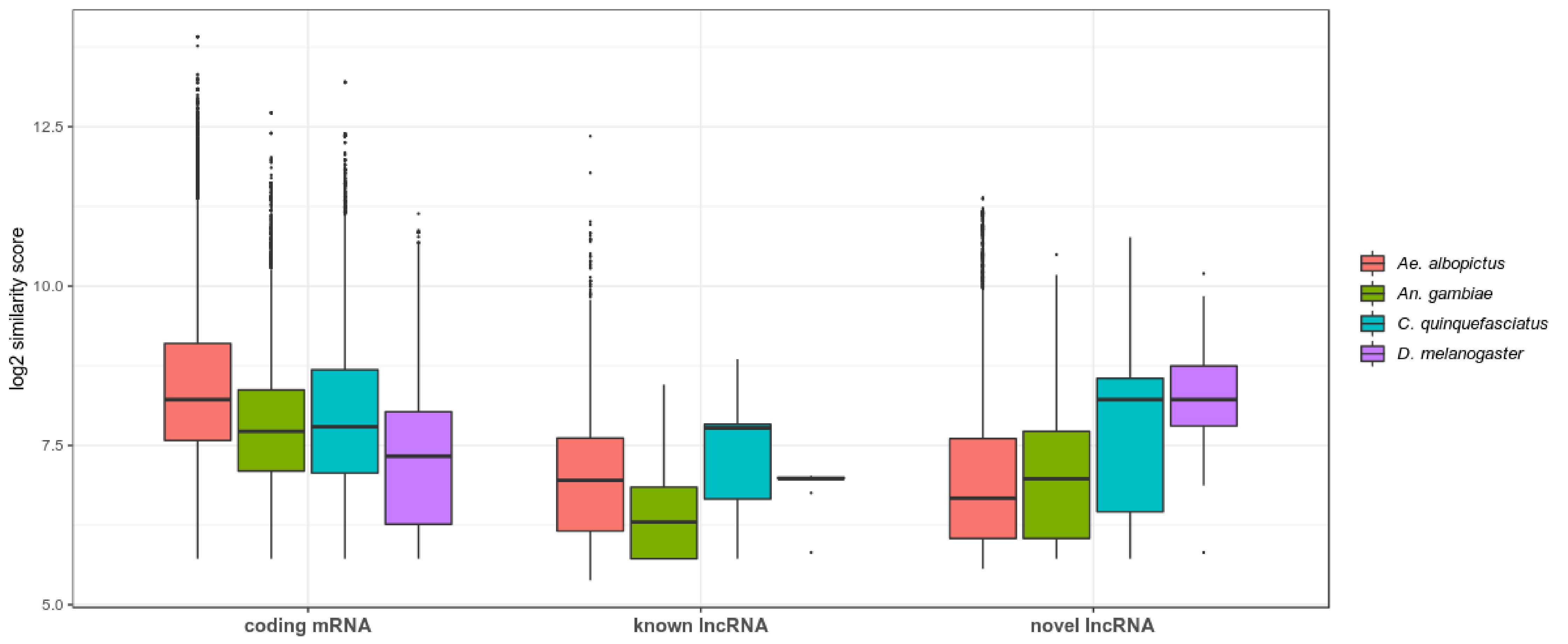

### Supplementary Figure S2

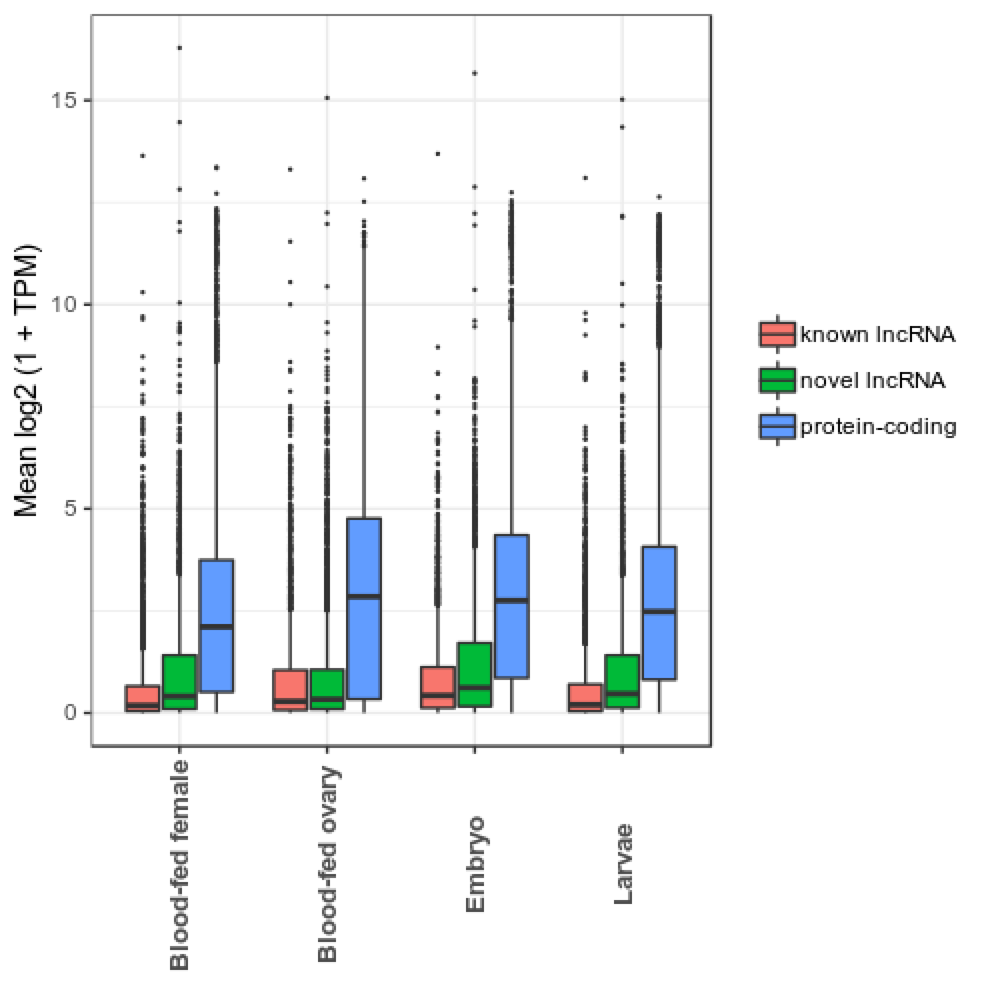

### Supplementary Figure S3

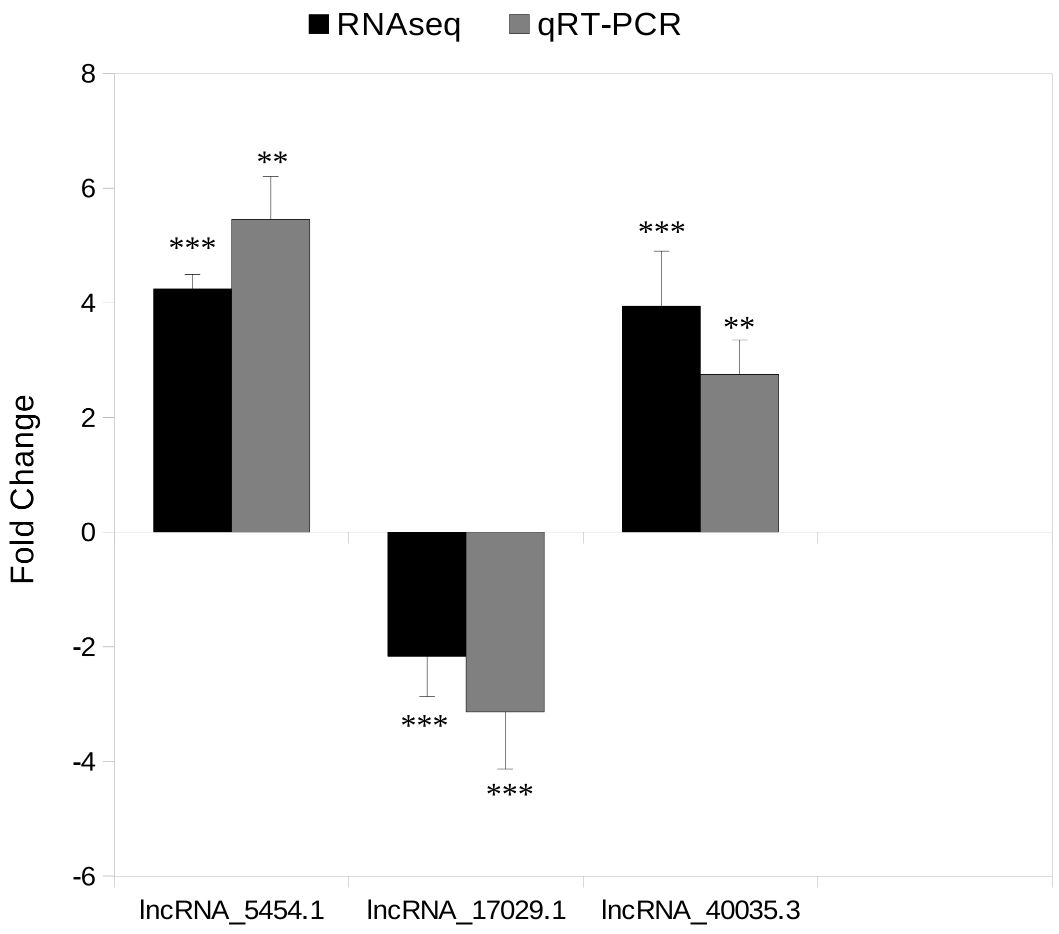
