## Supplementary Table S3 for "Systematic identification and characterization of *Aedes aegypti* long noncoding RNAs"

**S3 Table. List of primers used in this study.**

| **Primer** | **Sequence** | **Source** |
| --- | --- | --- |
| lncRNA_5454.1(Forward) | CCCTAACGCTCCTGACTCTG |  |
| lncRNA_5454.1 (Reverse) | ACACCTTTCAACCCAAGACG |  |
| lncRNA_17029.1 (Forward) | TGGTTGAGATCCACGACAAA |  |
| lncRNA_17029.1 (Reverse) | TTGCCGCAATGATTCAATAG |  |
| lncRNA_40035.3 (Forward) | TTGGGGATGCTACAAAGGTC |  |
| lncRNA_40035.3 (Reverse) | CACGGACGCATTACCAATTA |  |
| DENV1 (Forward) | CAAAAGGAAGTCGTGCAATA | Johnson et al. (2015) |
| DENV1 (Reverse) | CTGAGTGAATTCTCTCTACTGAACC |  |
| RPS17 (Forward) | AAGAAGTGGCCATCATTCCA | Dzaki et al. (2017) |
| RPS17 (Reverse) | GGTCTCCGGGTCGACTTC |  |
